# Zero-shot retention time prediction for unseen post-translational modifications with molecular structure encodings

**DOI:** 10.1101/2024.12.18.629045

**Authors:** Ceder Dens, Darien Yeung, Oleg Krokhin, Kris Laukens, Wout Bittremieux

## Abstract

Mass spectrometry-based proteomics relies on accurate peptide property prediction models to enhance peptide identification and characterization, especially when dealing with peptidoforms. However, current approaches are limited in their ability to generalize to peptides with novel post-translational modifications (PTMs) due to insufficient training data. To address this challenge, we introduce MoSTERT (Molecular Structure Transformer Encoder for Retention Time prediction) and its enhanced variant, MoSTERT-2S, two transformer-based models designed for zero-shot prediction of retention times of peptides with unseen PTMs. Unlike conventional models, MoSTERT encodes peptide residues at the molecular structure level, allowing it to handle diverse PTMs. MoSTERT-2S further improves accuracy by employing a two-step strategy: first predicting the retention time of the unmodified peptide, then estimating the retention time shift induced by the PTMs.

Evaluation on an external dataset demonstrates that MoSTERT-2S achieves state-of-the-art performance, reducing prediction errors compared to existing methods. Its ability to accurately predict retention times for peptides with a wide variety of PTMs not seen during training highlights its potential for advancing proteomic workflows analyzing proteoforms and protein modifications.

## Background

In recent years, the field of mass spectrometry (MS)-based proteomics has witnessed a series of significant breakthroughs, transforming our understanding of biological systems at the molecular level. These advancements encompass a wide array of achievements: from high-throughput techniques that enable the large-scale analysis of proteins during health and disease (1,2), over exploring the role of proteins across individual cells (3,4), to sophisticated methods for probing the dynamics of protein interactions (5,6).

A pivotal frontier in this evolving landscape is the detailed detection and characterization of various proteoforms. A prime example is the Human Proteoform Project, which aims to map the comprehensive array of protein forms arising from genetic variation, alternative splicing, and post-translational modifications (PTMs) (7). The identification of proteoforms, especially those carrying diverse PTMs, poses a formidable challenge, primarily due to the intricacies associated with PTM analysis and identification.

PTMs play critical functional roles, ranging from the regulation of protein activity and interactions to impacting protein stability and localization (8,9). The complexity of PTMs lies in their molecular diversity and the subtle yet significant ways they alter protein properties and functions (10). This complexity is further compounded by the fact that among the vast array of PTMs, only a few have been extensively studied. The limited understanding of the myriad PTMs, their potential occurrence, and their influence on experimental properties, such as liquid chromatography (LC) behavior, collisional cross section (CCS) from ion mobility, and fragmentation patterns during tandem mass spectrometry (MS/MS), presents a significant hurdle (11).

In recent years, peptide property prediction has emerged as a pivotal tool, significantly enhancing peptide identification performance. Tools such as Prosit (12) use deep neural networks to predict fragment ion intensities and retention times (RTs), and have been shown to be instrumental in peptide–spectrum match (PSM) rescoring (13). These advancements, however, are primarily focused on unmodified peptides or those with a limited diversity in modifications. This is largely because only these peptidoforms have sufficient training data available to develop robust ML models (14), which requires millions of unique training samples. While there have been efforts to generate ground truth synthetic measurements for modified peptides (11,15), these are insufficient for the development of large-scale, comprehensive deep learning models.

This gap highlights an urgent need for innovative and powerful computational tools tailored for modified peptides. A few existing peptide property prediction tools attempt to address this challenge (16,17) but are constrained by their limited PTM representation and inability to generalize across diverse and heterogeneous PTMs.

In this work, we introduce a transformer-based (18) indexed RT (iRT) prediction model, designed specifically for zero-shot predictions. This means that it can predict the iRT of peptides with any PTM, including those not encountered during training.

## Results and discussion

Our model, named MoSTERT (Molecular Structure Transformer Encoder for peptide Retention Time predictions), encodes the residues of a peptide as their molecular structure instead of using a distinct token for each amino acid. This lower level encoding gives a detailed representation of the input molecules, agnostic to whether they are canonical or modified amino acids. In addition, using the molecular structure allows the model to make accurate predictions for peptides with novel PTMs. First, MoSTERT generates an embedding for each residue using the Molecule Attention Transformer (MAT) model (19). MAT is a molecule encoder based on a transformer encoder, specialized for molecule input by incorporating atom distances and adjacency information. MAT was pretrained on 2 million molecules sampled from the ZINC15 database (20). This results in a sequence of embeddings, these embeddings are then processed by a transformer encoder (21) that predicts the RT (Figure 1a).

**Figure 1.**
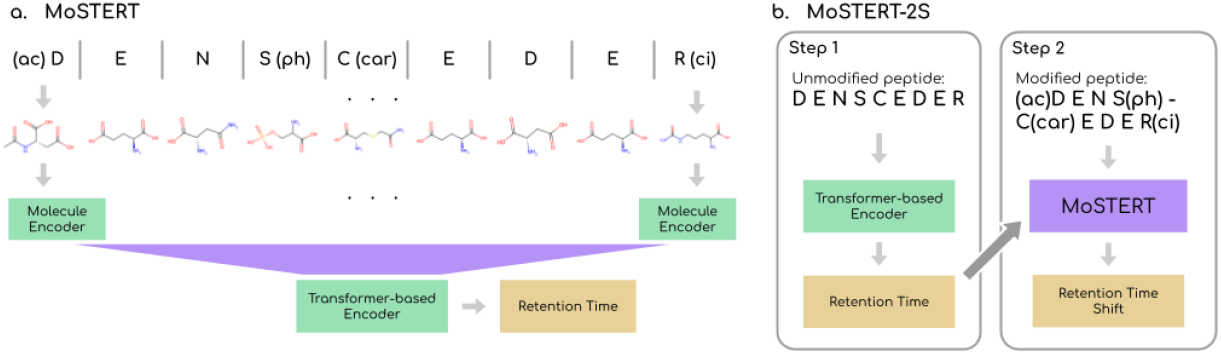
Model architectures. **(A)** Overview of the MoSTERT architecture. A modified peptide is first represented by a sequence of molecular structures. These structures are turned into a sequence of embeddings using a molecule encoder, MAT (19). The resulting embeddings are used as input to a transformer encoder, which is trained to predict the peptide’s iRT. **(B)** Overview of the architecture of MoSTERT-2S. In step one, the iRT of the unmodified version of the peptide is predicted with a transformer encoder that tokenizes the residues of the peptide. This prediction is added to the input of MoSTERT that is now trained to predict the RT shift between the unmodified and modified peptide.

To improve prediction accuracy, we introduce a second model using a two-step prediction approach (Figure 1b). This approach is motivated by the observation that RT prediction for unmodified peptides performs very well using sequence-based methods (12,22–24) and that abstractions at the level of amino acid token sequences have proven highly effective, as demonstrated by protein language models (25,26). Building on these insights, a transformer encoder is first used to predict the iRT of the tokenized unmodified peptide sequence, providing a robust baseline value. This baseline is then combined with the input to MoSTERT, which now predicts the iRT shift induced by the modification instead of the actual iRT. The iRT shift represents the difference between the predicted iRT of the unmodified peptide and the iRT of the modified peptide (27). By combining the powerful molecular representations of MoSTERT with the state-of-the-art performance of protein language models in a two-step framework, this strategy leverages the high prediction accuracy for unmodified peptides while enhancing MoSTERT’s superior ability to handle modifications. We refer to this model as MoSTERT-2S (MoSTERT-2-Step).

### A two-step approach improves zero-shot iRT predictions

We begin by comparing MoSTERT to MoSTERT-2S using the Chronologer dataset (28) and a leave-one-group-out cross-validation strategy (Figure 2). To prevent data leakage and accurately assess the models’ ability to predict peptides with unseen PTMs, we group related PTMs together. A set of the most common PTMs (carbamidomethylation, oxidation, acetylation, and phosphorylation) is consistently used for training, while the remaining five PTM groups are designated as the test set for each iteration of the leave-one-group-out cross-validation. All peptides in the test set contain PTM(s) that were not encountered during training and validation.

**Figure 2.**
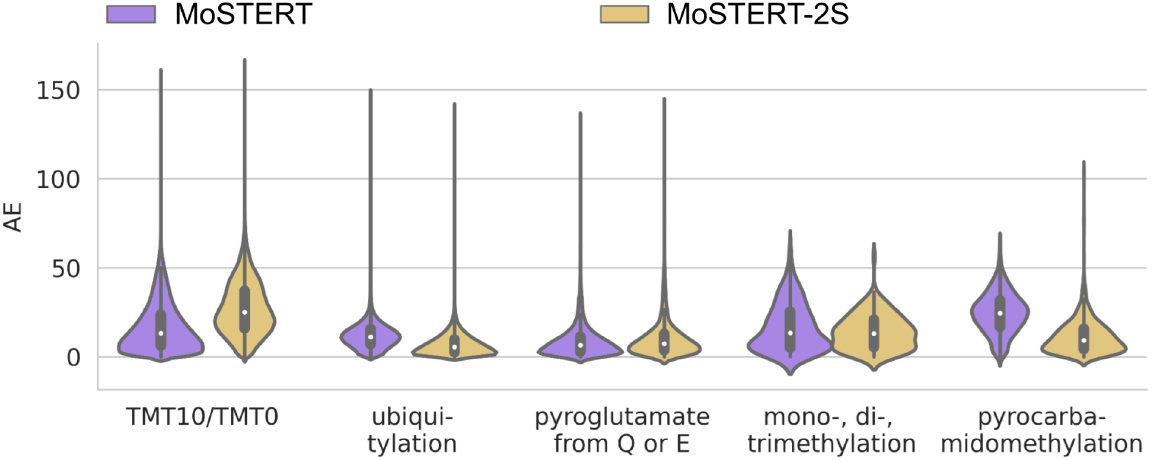
RT prediction performance on Chronologer data. Violin plots of the absolute error (AE) of the iRT predictions by MoSTERT and MoSTERT-2S, evaluated by a leave-one-group-out cross-validation strategy on the Chronologer dataset (28). In each test set, all peptides contain at least one PTM that was not seen during training.

This comparison shows that MoSTERT and MoSTERT-2S yield similar results for the pyroglutamate and methylation PTM groups (Figure 2). MoSTERT shows better performance on the TMT group, while MoSTERT-2S outperforms MoSTERT on ubiquitylation and pyrocarbamidomethylation. These findings indicate that the two-step strategy yields promising results for predicting retention times of peptides with unseen PTMs.

### Zero-shot accuracy is highly dependent on the PTM

For a more realistic comparison, we generate training, validation, and test splits that mimic the scenario where a substantial amount of training data is available but limited to a small set of PTMs. For evaluating the model’s zero-shot performance, the test set again consists exclusively of peptides containing at least one PTM not encountered during training. We also include DeepLC (16) in the evaluation. DeepLC is a deep learning model based on a convolutional neural network that encodes modified peptides using various features, including the atomic composition of amino acids and their modifications, also allowing predictions for peptides with novel PTMs.

The training dataset is derived from the Chronologer dataset (28), with all peptides containing a mono-, di-, or trimethylation removed, as these PTMs are abundant in the test dataset. This leaves 12 unique PTMs available for training and validation. To create training-validation splits, the set of the most common PTMs is always included in the training data, while the remaining PTMs are divided into four groups of related PTMs. Using a leave-one-group-out strategy, we generate four train-validation splits.

The test dataset consists of the ProteomeTools PTM dataset (11) and a dataset with dimethylated peptides generated by Neale et al. (27). Only peptides with at least one PTM not in the training dataset were retained. This results in a test dataset containing 15 unique PTMs.

We calculate the absolute error (AE) of the predicted iRT for each test peptide and group the results by PTM. We find that the prediction performance varies depending on the PTM. Certain PTMs, such as dimethylation, biotinylation, and nitrotyrosine, prove challenging to predict for all models, while others, like monomethylation and glutarylation, are much easier to predict (Figure 3). We hypothesize that this variability may be due to factors like the structural similarity of certain PTMs to those in the training data or unique physicochemical properties of certain PTMs, causing them to behave differently than expected.

**Figure 3.**
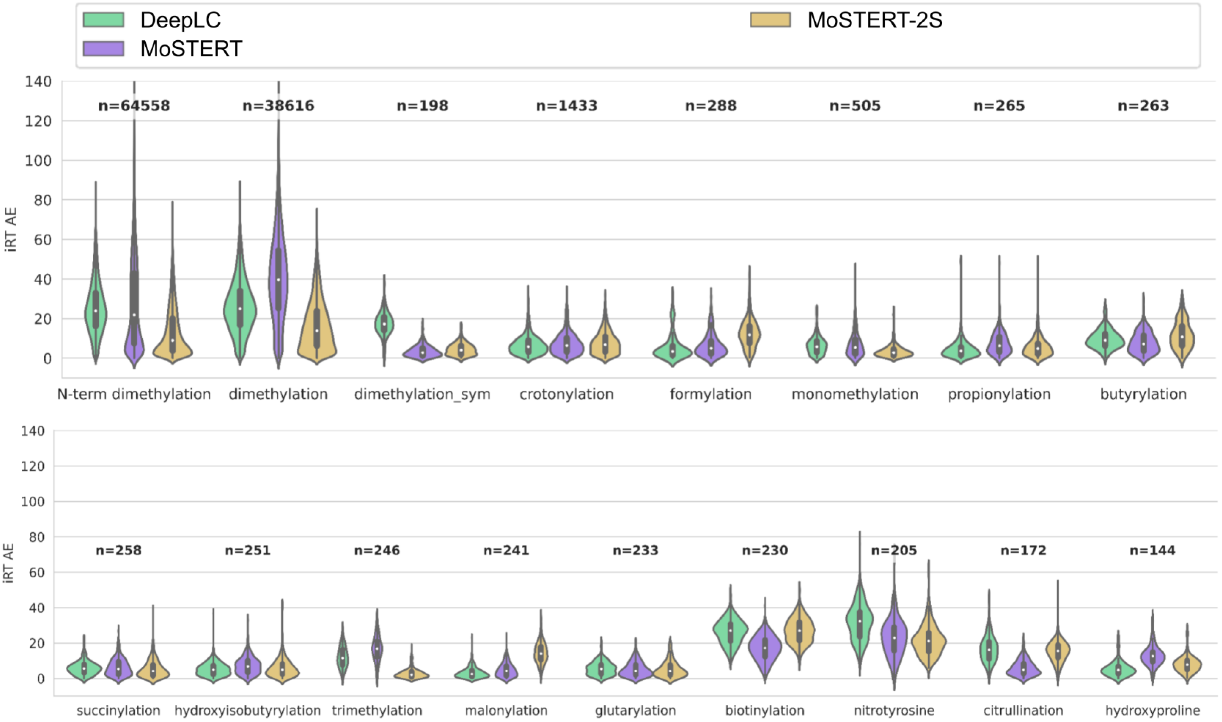
Absolute error (AE) of the predicted iRT per PTM. Violin plots showing the prediction errors of DeepLC (16), MoSTERT, and MoSTERT-2S across all PTMs, none of which were seen during training. The prediction for peptides containing multiple unseen PTMs will contribute to the results of each PTM. The number of test peptides for each PTM is given above the corresponding violin plots.

### MoSTERT-2S achieves the highest retention time prediction performance on novel PTMs

To estimate and compare the overall iRT prediction performance for peptides with unseen PTMs, we evaluate DeepLC, MoSTERT, and MoSTERT-2S using two metrics: the AE across all test peptides (micro AE) and the mean AE (MAE) per PTM (macro PTM MAE).

MoSTERT shows a slightly lower median micro AE than DeepLC but has a larger variability. While many of MoSTERT’s predictions have a lower AE, it also has slightly more peptides with larger errors (Figure 4a). In contrast, MoSTERT-2S clearly outperforms both DeepLC and MoSTERT, showing a substantially lower median AE and a narrower AE distribution overall.

**Figure 4.**
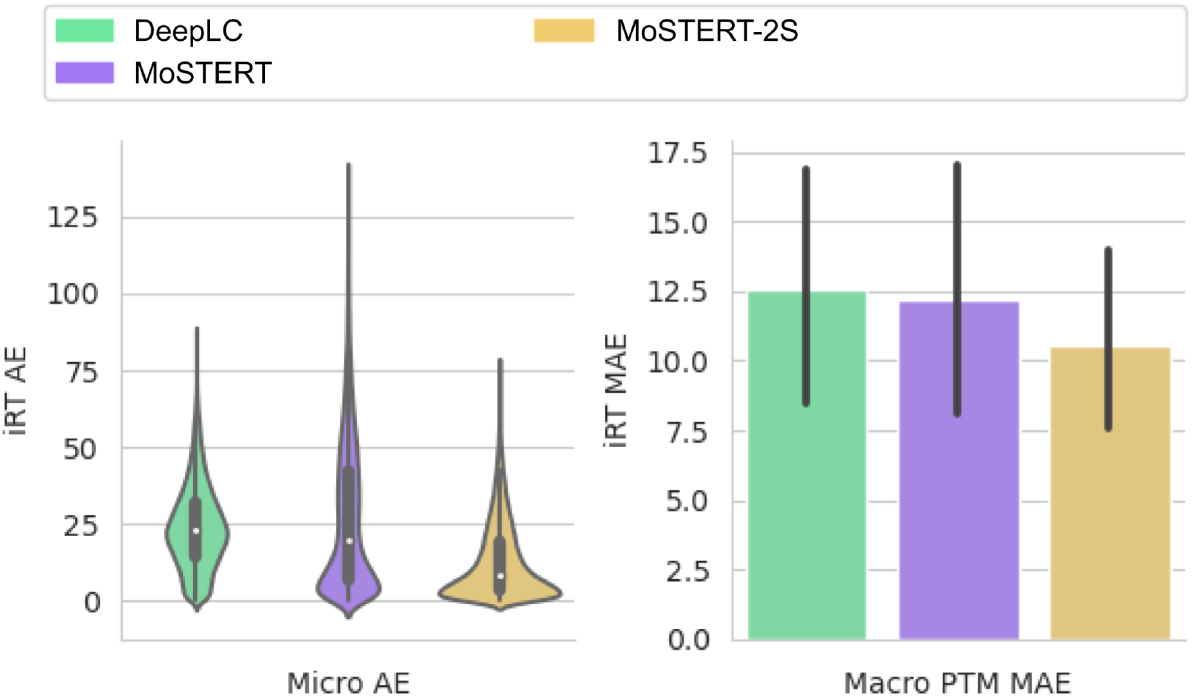
Micro AE and macro PTM MAE of iRT predictions. **(A)** Violin plots show the AE distribution for all test peptides (micro AE) predicted by DeepLC, MoSTERT, and MoSTERT-2S. Boxplots show the quartiles of the data with whiskers extending to 1.5^*^IQR. **(B)** The MAE of the iRT predictions were calculated per PTM. The bars represent the mean of these 15 PTM MAE’s and the error bars show ± the standard deviation. DeepLC achieves an average PTM MAE of 12.54 ± 9.17, MoSTERT has an average of 12.18 ± 9.71, and MoSTERT-2S shows the best performance with an average PTM MAE of 10.57 ± 6.57.

The PTM-specific MAE shows a similar trend (Figure 4b): MoSTERT has a slightly lower average PTM MAE than DeepLC and MoSTERT-2S clearly shows the best performance.

## Conclusions

This work introduces a novel approach for predicting the iRT of peptides with previously unseen PTMs, aiming to address the challenges related to identifying a high diversity of proteoforms. MoSTERT and MoSTERT-2S, our two proposed transformer-based architectures, demonstrate promising advancements in handling modified peptides with PTMs unseen during training. Both models capture peptide structure by encoding the molecular representations of amino acids and PTMs. MoSTERT-2S leverages a two-step prediction strategy yielding improved prediction accuracy for zero-shot RT predictions of peptides with novel PTMs compared to state-of-the-art methods. This is a critical step towards a broader applicability in proteomics workflows where the lack of sufficient training data for a broad range of PTMs has previously limited computational prediction models.

## Methods

### MoSTERT and MoSTERT-2S

We developed two molecular structure transformer encoder (18,21) models: MoSTERT and MoSTERT-2S. MoSTERT takes a (potentially modified) peptide as input and directly predicts the iRT, while MoSTERT-2S uses a two-step process. In MoSTERT-2S, a transformer encoder first predicts the iRT of the unmodified peptide, which then serves as an additional input to MoSTERT’s architecture, now trained to predict the iRT shift caused by the modified peptidoform.

The input to MoSTERT consists of a peptide sequence that can contain one or more PTMs. Each PTM is defined through a SMARTS reaction (29) (Table S1), which describes the molecular change a canonical amino acid undergoes when it is modified by the PTM. For peptides with unseen PTMs, we simply add a SMARTS reaction that applies the modification on the relevant amino acid. The molecular structures of amino acids used are provided by RDKit (30) and RDKit also applies the SMARTS reactions. This generates a molecular representation for each residue, including any PTMs.

To embed these molecular representations, we use the Molecule Attention Transformer (MAT) (19), a pretrained model designed for molecular property prediction. MAT is based on a specialized transformer encoder and incorporates atom distances and adjacency information. MAT was pretrained on 2 million molecules sampled from the ZINC15 database (20). MAT generates a molecular embedding for each molecular structure. The sequence of these embeddings are used as input to a transformer encoder from TAPE (31), initialized with random parameters. The output embeddings are processed through two fully connected layers with dimensions [embedding size, ⅔ ^*^ embedding size] and [⅔ ^*^ embedding size, 1], outputting the predicted iRT (see Table 1 for embedding sizes). The model is optimized using the MAE as the loss function.

**Table 1.** Hyperparameters selected for MoSTERT and MoSTERT-2S. This table presents the chosen hyperparameters for the single-step model (MoSTERT) and for both steps in the two-step model (MoSTERT-2S). Various configurations were tested for learning rate, number of transformer layers, and embedding size.

|  | Learning rate | Number of layers | Embedding size |
| --- | --- | --- | --- |
| MoSTERT | 1e-4 | 8 | 384 |
| MoSTERT-2S step 1 | 3e-5 | 8 | 96 |
| MoSTERT-2S step 2 | 1e-5 | 4 | 384 |

MoSTERT-2S uses a two-step prediction strategy. In the first step, a transformer encoder predicts the iRT of the unmodified version of the peptide sequence. This model, based on TAPE, tokenizes the peptide sequence and was first pretrained using masked language modeling before being fine-tuned on the training dataset (32).

In the second step, a variation of MoSTERT incorporates the predicted iRT of the unmodified peptide. This value is converted into a vector of the same embedding size via a fully connected layer and then added to the sequence of molecular embeddings. This modified MoSTERT model is trained to predict the iRT shift caused by the PTMs, defined as the difference between the actual iRT of the modified peptide and the predicted iRT of the unmodified peptide. The final iRT prediction for the modified peptide is calculated by adding the predicted iRT shift to the initial unmodified iRT prediction.

The hyperparameters for MoSTERT and MoSTERT-2S were manually tuned by adjusting the number of transformer layers (2, 4, 8, or 12), embedding size (48, 96, 192, 384, or 768) and learning rate (1e-6, 3e-6, 1e-5, 3e-5, 1e-4, 3e-4, or 1e-3) (Table 1). Additionally, each model was trained with a batch size of 1024, using the Adam optimizer (33). A warmup phase was added to the learning rate, increasing it from 1% to 100% over the first 10 epochs. Early stopping based on the validation dataset was used to determine when the model reached convergence.

### DeepLC

DeepLC (16) is a peptide RT prediction model based on a convolutional neural network architecture. As input, it takes a one-hot encoding of the unmodified peptide sequence, along with the atom counts of C, H, N, O, P, and S of each amino acid in the peptide sequence. This allows DeepLC to make RT predictions for peptides with PTMs not seen during training. For consistency, DeepLC was always retrained and evaluated using the same train-test splits as our models. However, DeepLC internally creates its own train-validation splits.

### Data

The evaluations were done using data from three sources: the dataset collected for Chronologer (28), the ProteomeTools PTM dataset (11), and data generated by Q. Neale et al. (27).

#### Leave-one-group-out cross-validation on the Chronologer dataset

For the leave-one-group-out cross-validation, only the Chronologer dataset (28) was used. This dataset consists of more than 2.6 million peptides from various sources, containing tryptic peptides, immunopeptides, as well as peptides originating from multiple organisms. This dataset features peptides with 13 different PTMs, present on various amino acids. Due to potential variations from different experimental conditions, such as different instruments, different laboratories, and experiments performed on different days, the RT has to be standardized to iRT. This calibration process was already applied for the Chronologer data set.

To further ensure data integrity, quality, and consistency, several data curation steps were undertaken. First, all peptides with an iRT of more than 150 were deemed anomalous as they fall outside of the chromatographic range and were removed, eliminating around 132,000 peptides. Subsequently, we found recurring identical iRT values across numerous distinct peptides. These stem from technical or postprocessing artifacts, and we removed all peptides associated with iRT values appearing 300 times or more from the same source, resulting in the removal of around 35,000 peptides. Additionally, peptides featuring PTMs deemed impossible, such as M acetylation and Q acetylation, were discarded, amounting to approximately 1400 peptides. Lastly, duplicate peptides were merged by computing their median iRT values, resulting in the loss of around 367,000 peptides. This curation process resulted in an iRT dataset comprising 2.1 million unique peptides containing 12 PTMs. Note that the curation process retained none of the peptides with a succinylation modification.

To create cross-validation splits for this evaluation, we grouped related PTMs to prevent data leakage and to accurately evaluate unseen PTM prediction performance. The first group includes TMT0 and TMT10, occurring at the N-terminus or on lysine (K). The second group consists of pyroglutamate from glutamic acid (E) and from glutamine (Q). The third group includes monomethylation of K and arginine (R), asymmetric dimethylation of K and R, and trimethylation of K. Groups 4 and 5 contain only one PTM each: ubiquitylation of K and pyrocarbamidomethylation of cysteine (C), respectively. The remaining PTMs, including carbamidomethylation of C, methionine (M) oxidation, acetylation of K, N-terminal acetylation, and phosphorylation of serine (S), threonine (T), and tyrosine (Y), are always included in the training dataset.

In the leave-one-group-out cross-validation setup, each of these five PTM groups is used as a test set while the remaining four groups are added to the training and validation dataset. Thus, each peptide in the test set will always contain a PTM that is never seen during training or validation. To avoid data leakage, peptides with the same unmodified sequence as those in the test set are removed from the training and validation dataset. For each cross-validation split, the training and validation dataset is further divided using a similar strategy. Within each split, the remaining four PTM groups are rotated, with one group serving as validation data and the other three groups as training data. Again, peptides in the training set with unmodified sequences identical to those in the validation data are removed, ensuring complete separation of test, validation, and training sequences across all splits.

#### External test dataset split

We generated training, validation, and test splits to better represent a realistic scenario, where a substantial amount of training data is available for only a limited number of PTMs. The training and validation dataset is derived from the Chronologer dataset (28), preprocessed similarly to the leave-one-group-out cross-validation described in the previous section. Additionally, all peptides with mono-, di-, and trimethylation were excluded, as these PTMs are abundant in our test dataset. While DeepLC (16) creates its own training-validation splits, our model’s splits were created again by using a fixed set of PTMs that are always included in the training data: carbamidomethylation of C, M oxidation, acetylation of K, N-terminal acetylation, and phosphorylation of S, T, and Y. The remaining PTMs were divided into the same groups as in the previous section, excluding the methylation group. This approach generated four training-validation splits using a leave-one-group-out strategy. Peptides in the training dataset with the same unmodified sequence as those in the validation dataset were excluded.

The test dataset consists of the Proteometools PTM (11) dataset and data for dimethylated peptides generated by Q. Neale et al. (27). To process the ProteomeTools PTM data, we read the PTM-combined evidence.txt files and filtered for peptides with an Andromeda score above 90 and a posterior error probability (PEP) below 1%. Peptides with only one measurement were removed and duplicate peptides were merged by calculating their median RT. The RT measurements of the same peptide differed significantly between data files in some cases. Therefore, we calculated the iRT values by linearly calibrating RTs per file using the standardized RT of reference peptides (34). We also included a dataset of dimethylated peptides, both N-terminal and on K. The RTs of this dataset were calibrated using the peptides overlapping with the training-validation dataset.

Only peptides with a PTM not in the training-validation dataset were retained, resulting in a zero-shot test dataset containing 16 unique PTMs. For this dataset, results are presented separately for N-terminal dimethylation, and dimethylation of K. Peptides in the training-validation dataset with the same unmodified sequence as a peptide in the test dataset were again removed.

### Consensus strategy

In addition to the two-step prediction strategy, MoSTERT-2S employs a consensus prediction strategy to minimize the effect of specific training-validation splits on the final prediction performance. The training-validation dataset has a limited PTM diversity and we want to simulate a zero-shot scenario during validation. This means that the selection of PTMs for the validation dataset could have an impact on the predictions. To mitigate this effect, we create multiple training-validation splits and train a model on each one. For the prediction on test peptides, we calculate the median of the predicted iRT values across all trained models.

## Acknowledgements

This work was supported by the Flemish Government (Flanders AI Research Program) and the University of Antwerp Industrial Research Fund. The computational resources (Stevin Supercomputer Infrastructure) and services used in this work were provided by the VSC (Flemish Supercomputer Center), funded by Ghent University, FWO, and the Flemish Government – department EWI.

## Supplementary material

**Table S1.**
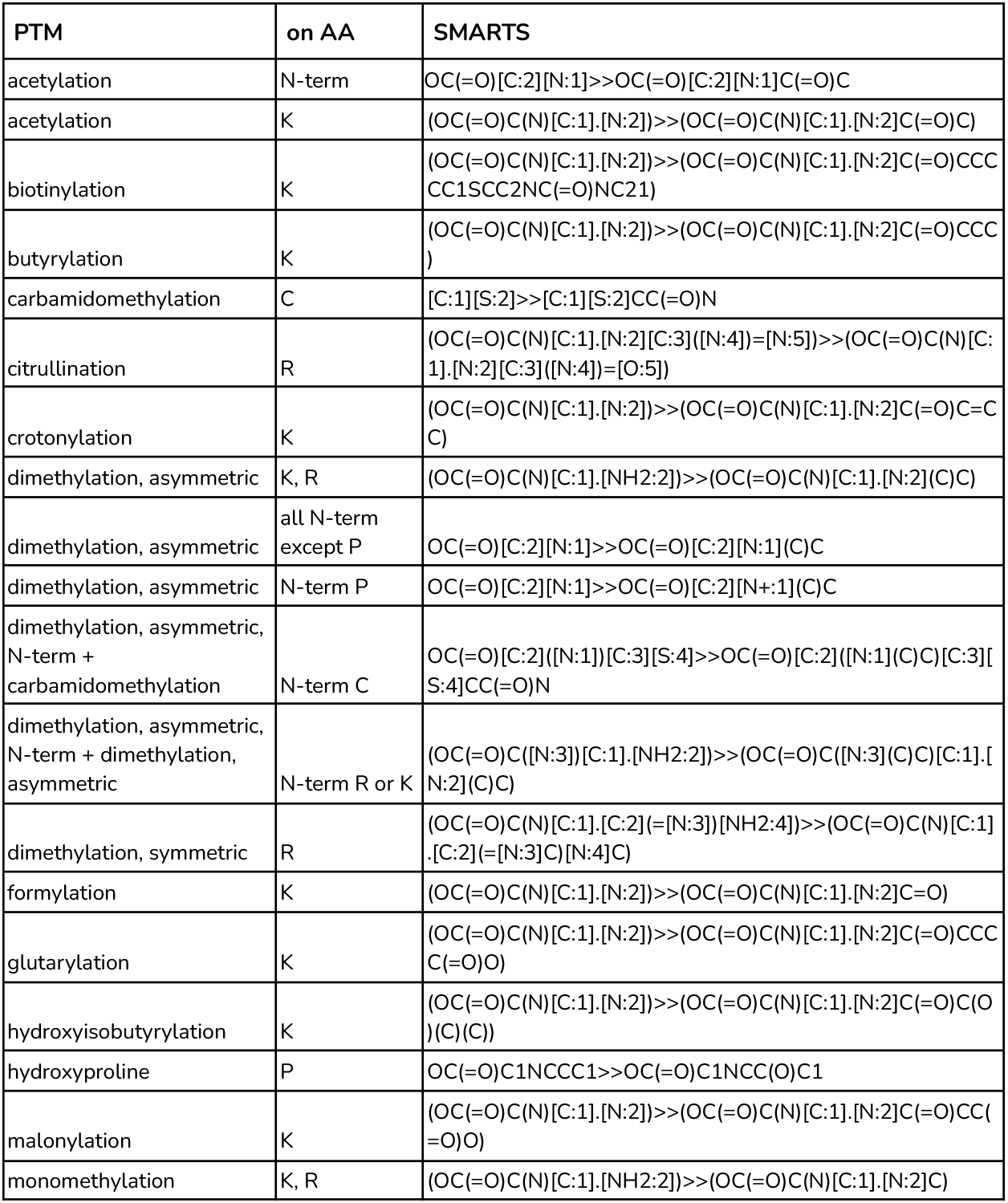
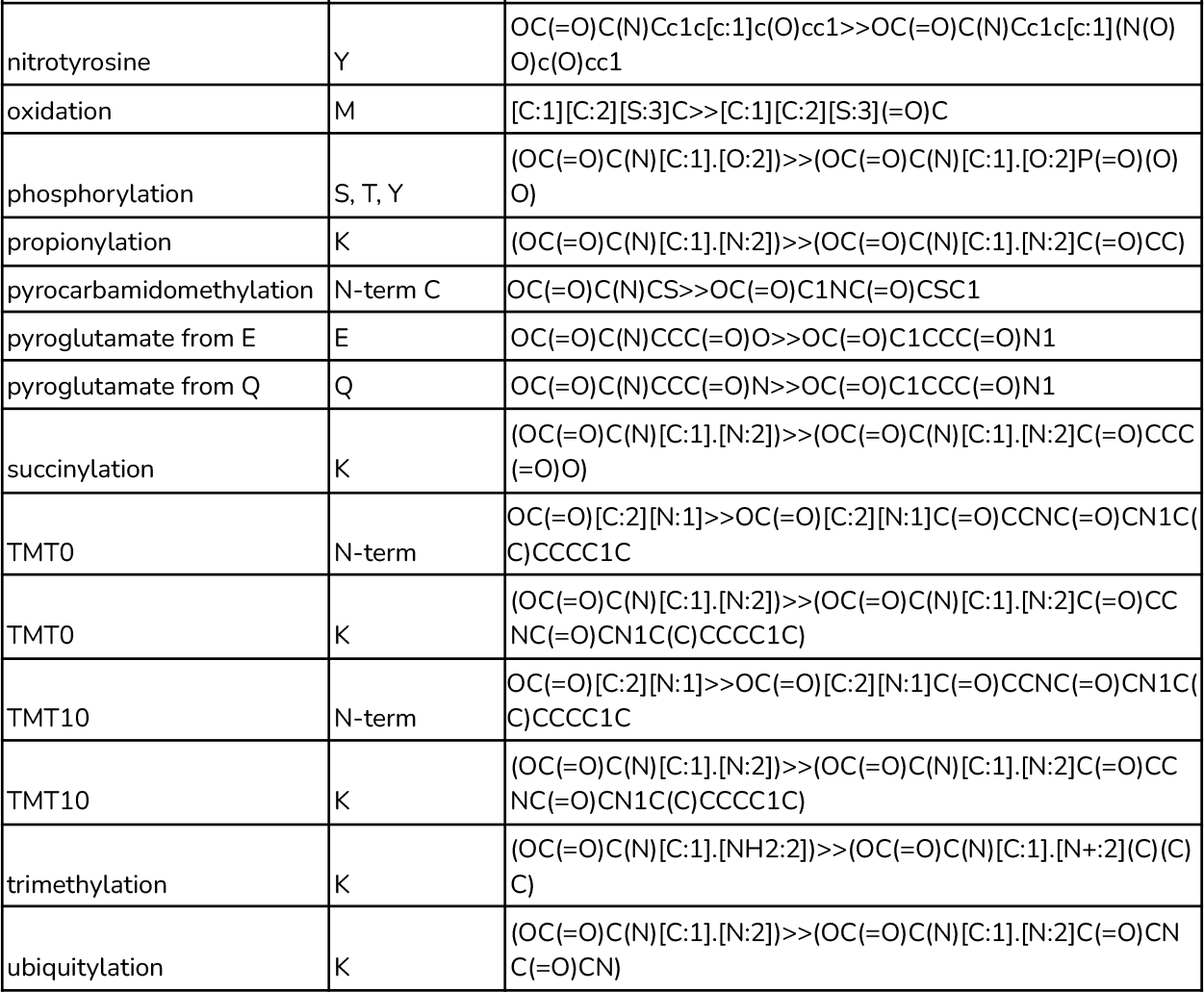
SMARTS reactions. SMARTS reactions for the PTMs present in the datasets used in this study and on which amino acid they will be applied. SMARTS reactions are usually different when the PTM has to be applied on the N-terminus or on the side chain. A few reactions combine an N-terminal and a side-chain PTM. SMARTS reactions are only guaranteed to work with the given amino acids.

